# Association between substance use disorder and polygenic liability to schizophrenia

**DOI:** 10.1101/129288

**Authors:** Sarah M. Hartz, Amy Horton, Mary Oehlert, Caitlin E. Carey, Arpana Agrawal, Ryan Bogdan, Li-Shiun Chen, Dana B. Hancock, Eric O. Johnson, Carlos Pato, Michele Pato, John P. Rice, Laura J. Bierut

## Abstract

**Background:** There are high levels of comorbidity between schizophrenia and substance use disorder, but little is known about the genetic etiology of this comorbidity.

**Methods:** Here, we test the hypothesis that shared genetic liability contributes to the high rates of comorbidity between schizophrenia and substance use disorder. To do this, polygenic risk scores for schizophrenia derived from a large meta-analysis by the Psychiatric Genomics Consortium were computed in three substance use disorder datasets: COGEND (ascertained for nicotine dependence n=918 cases, 988 controls), COGA (ascertained for alcohol dependence n=643 cases, 384 controls), and FSCD (ascertained for cocaine dependence n=210 cases, 317 controls). Phenotypes were harmonized across the three datasets and standardized analyses were performed. Genome-wide genotypes were imputed to 1000 Genomes reference panel.

**Results:** In each individual dataset and in the mega-analysis, strong associations were observed between any substance use disorder diagnosis and the polygenic risk score for schizophrenia (mega-analysis pseudo R^2^ range 0.8%-3.7%, minimum p=4×10^-23^).

**Conclusions:** These results suggest that comorbidity between schizophrenia and substance use disorder is partially attributable to shared polygenic liability. This shared liability is most consistent with a general risk for substance use disorder rather than specific risks for individual substance use disorders and adds to increasing evidence of a blurred boundary between schizophrenia and substance use disorder.

## Introduction

Schizophrenia and substance use disorder frequently co-occur in the same individual (1-6). This increased comorbidity can be explained through several, non-exclusive mechanisms (7) (Figure 1): 1. schizophrenia may cause the development of substance use disorder (8); 2. substance use disorder may lead to the onset of schizophrenia (9); or 3. there may be common underlying risk factors, environmental and genetic, that predispose to both schizophrenia and substance use disorder (10, 11). With the publication of large meta-analyses of genome-wide association studies (GWAS), polygenic risk scores now can be used to measure the shared genetic liability between schizophrenia and substance use disorder, which can lead to better understanding of potential mechanisms for these comorbid conditions.

**Figure 1:**
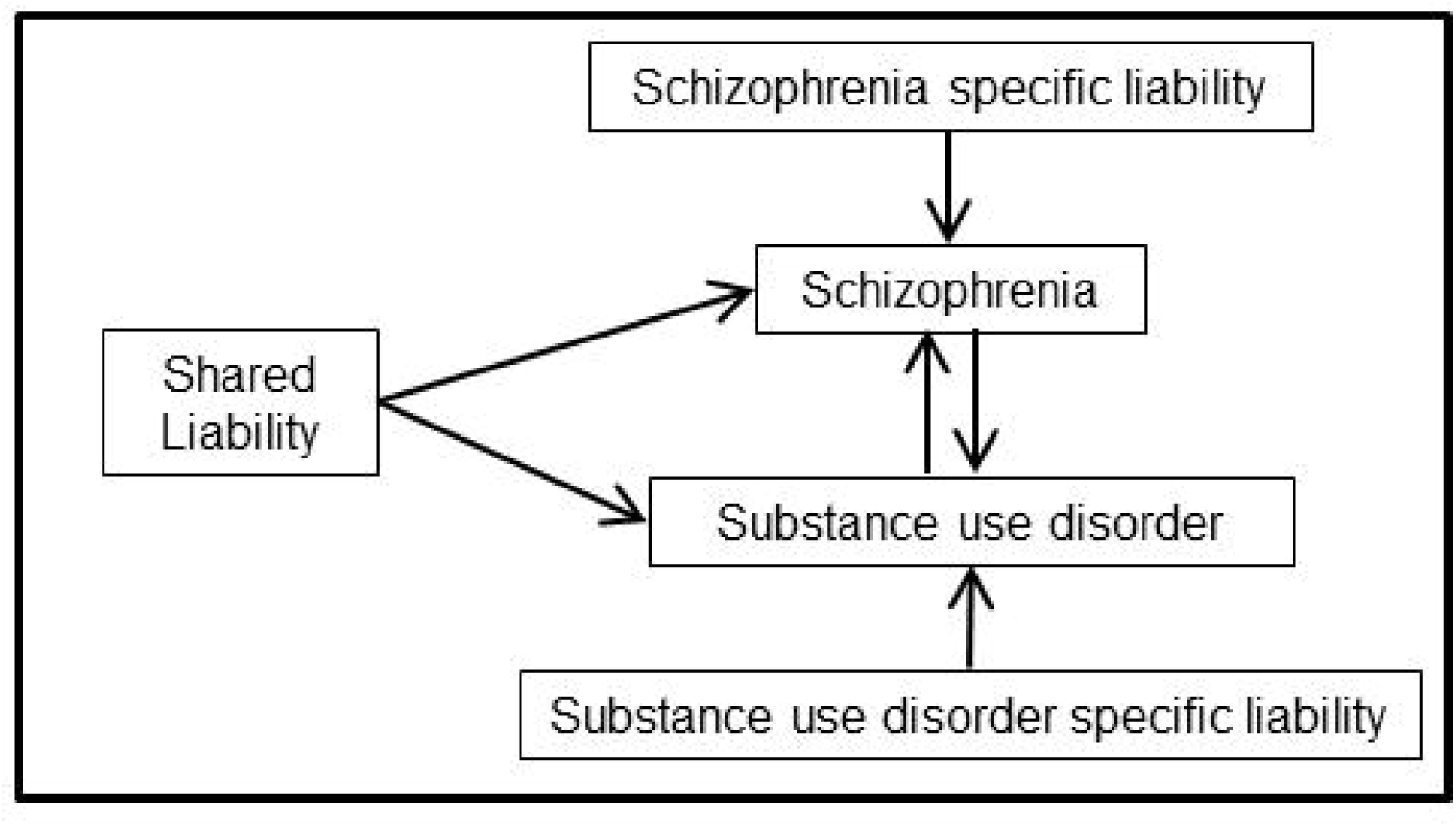
Model of liability leading to the comorbidity between schizophrenia and substance dependence.

Polygenic risk scores represent aggregated effects across the many loci associated with a disorder at p-value thresholds that accommodate tens of thousands of SNPs, thus approximating additive genetic variance (12). Polygenic risk scores are generated using a discovery genetic association study of one disorder (e.g. schizophrenia meta-analysis) and can be applied to compute the phenotypic variance explained by the score in a new independent sample. For example, polygenic risk scores were used to show that schizophrenia has underlying shared genetic liability with bipolar disorder (12-17) and major depressive disorder (18). Importantly, a growing number of studies have begun to investigate shared genetic liability between schizophrenia and patterns of substance use. We recently found a statistically significant association between general liability for substance use disorder and polygenic risk for cross-disorder psychopathology (19). In addition, recent studies have described common genetic risk factors between schizophrenia and cannabis use (20, 21), and evidence for shared genetic factors between schizophrenia and smoking-related phenotypes (22, 23).

Despite recent progress, to our knowledge there has been no comprehensive and systematic examination of the association between polygenic liability to schizophrenia and substance use disorder, which has been carefully assessed for licit and illicit substances. For these analyses, we leverage studies systematically ascertained for substance use disorder to determine whether the schizophrenia polygenic risk scores are associated with these substance dependences.

## Methods

### Datasets

Three datasets were used for these analyses (Table 1): the Collaborative Genetic Study of Nicotine Dependence (24-28) (COGEND) was ascertained for nicotine dependence; the Collaborative Study on the Genetics of Alcoholism (29-33) (COGA) was ascertained for alcohol dependence; and the Family Study of Cocaine Dependence (34) (FSCD) was ascertained for cocaine dependence. Individuals from each of the three datasets were used to comprise the Study of Addiction: Genetics and Environment (SAGE) (35) (dbGaP accession number phs000092.v1.p1). Additional participants from the COGEND study were subsequently added to the SAGE study (dbGaP accession number phs000404.v1.p1). For this study, we restricted analyses to self-reported non-Hispanic individuals of European descent (N=3,676) because this is the population used to derive the polygenic risk score for schizophrenia. Ancestry was confirmed through principal component analyses. All studies were approved by local Institutional Review Boards, and all participants provided informed consent.

**Table 1:** Participant Characteristics by Study

| Study | COGEND |  | COGA |  | FSCD |  |
| --- | --- | --- | --- | --- | --- | --- |
| Ascertainment | Nicotine Dependence |  | Alcohol Dependence |  | Cocaine Dependence |  |
|  | Cases | Controls | Cases | Controls | Cases | Controls |
| N | 918 | 988 | 801 | 442 | 210 | 317 |
| % Female | 53% | 69% | 30% | 74% | 48% | 52% |
| Average Age (SD) | 37 (5) | 36 (5) | 43 (11) | 50 (12) | 33 (9) | 34 (9) |
| % Tobacco Use Disorder | 100% | 0% | 71% | 28% | 71% | 8% |
| % Alcohol Use Disorder | 18% | 7% | 100% | 0% | 100% | 22% |
| % Cannabis Use Disorder | 13% | 4% | 29% | 0% | 7% | 63% |
| % Stimulant Use Disorder | 11% | 3% | 38% | 0% | 100% | 0% |

## Recruitment

### COGEND

The Collaborative Genetic Study of Nicotine Dependence (COGEND) was initiated to detect and characterize genes that alter risk for tobacco use disorder. Community-based recruitment enrolled nicotine dependent cases and non-dependent smoking controls in St. Louis, Missouri and Detroit, Michigan between 2002 and 2007. All participants were between the ages of 25-44 years and spoke English. Nicotine dependent cases were defined as current smokers with a Fagerström Test for Nicotine Dependence (FTND) score of 4 or greater(24). Control status was defined as smoking at least 100 cigarettes lifetime, but never being nicotine dependent (lifetime FTND score < 1). Other substance use disorder diagnoses or comorbid disorders were not used as exclusionary criteria.

### COGA

The Collaborative Study on the Genetics of Alcoholism (COGA), initiated in 1989, is a large-scale family study with its primary aim being the identification of genes that contribute to alcoholism susceptibility. Participants were recruited from 7 sites across the U.S. Alcohol dependent probands were recruited from treatment facilities. Family members of the alcohol dependent probands were recruited and comparison families were drawn from the same communities. An alcohol dependent case-control sample of biologically unrelated individuals was drawn from COGA subjects (dbGaP accession number phs000125.v1.p1). All cases met DSM-IV criteria for alcohol dependence, and controls were defined as individuals who consumed alcohol, but did not meet any definition of alcohol dependence or alcohol abuse, nor did they meet any DSM-IIIR or DSM-IV definition of abuse or dependence for other drugs (except nicotine).

### FSCD

The Family Study of Cocaine Dependence (FSCD) was initiated in 2000 with the primary goal of increasing understanding of the familial and non-familial antecedents and consequences of stimulant use disorder(36). Individuals with cocaine dependence defined by DSM-IV criteria were systematically recruited from chemical dependency treatment units in the greater St. Louis metropolitan area. Community-based control participants were identified and matched by age, race, gender, and residential zip code.

### Assessments

All participants were assessed for baseline demographics and a comprehensive history of substance use and problem use. The Semi-Structured Assessment for the Genetics of Alcoholism (SSAGA) (30, 37), a validated instrument developed by COGA which provides a detailed evaluation of alcohol, tobacco, other substance use disorder, and psychiatric disorders, was the foundation of all assessments. Because a similar assessment protocol was used across all studies, all phenotypes were easily harmonized across datasets. Tobacco use disorder was defined as a score of 4 or greater on the FTND (24) when smoking the most. Alcohol use disorder, stimulant use disorder, and cannabis use disorder were defined by lifetime history of DSM-IV dependence criteria (38). The phenotype of any substance use disorder was defined as meeting the above criteria for nicotine dependence, alcohol dependence, cocaine dependence or marijuana dependence.

### Genotyping

A common analytic pipeline was used to process and impute genotypes across all studies. COGEND samples were genotyped on either the Illumina Human1M (dbGaP accession number phs000092.v1.p1) or the Illumina 2.5M (as part of dbGaP accession number phs000404.v1.p1) platforms. These datasets were combined and genotype data from the intersection of the 1M and 2.5M platforms were used as the basis for imputation (28). COGA and FSCD participants were genotyped on the Illumina Human 1M platform. A standardized procedure was used to impute the three studies. All samples were imputed using IMPUTE2(39, 40) with 1000 Genomes Phase 3 (Oct. 2014 release) (41) as the reference panel. Variants with info score <0.3 were excluded; variants with minor allele frequency (MAF) < 0.02 were excluded; and genotypes with probabilities <0.9 were treated as missing genotypes. Hard call genotypes were then constructed for the polygenic risk score analyses.

### Statistical Analyses

The results from the Psychiatric Genomics Consortium (PGC) schizophrenia GWAS meta-analysis (PGC-SCZ2, N=74,626) (42) were used to generate schizophrenia polygenic risk scores for participants in the three independent datasets using the genotype dosages from the imputed data. There is no overlap of participants between these three datasets and the PGG-SZ2 study. We used the summary statistics from the PGC-SCZ2 European case control samples for variants with imputation info >=0.9 and MAF >= 0.02. Schizophrenia polygenic risk scores were calculated for a series of p-value thresholds (from 1×10^-5^ to 0.5) in PGC-SCZ2 using a modified version of PRSice (43), an R language (44) wrapper script using second generation PLINK (27, 45). SNPs in each dataset were pruned by PRSice (version 1.23) (43) using p-value-informed linkage disequilibrium (LD) clumping: *R^2^* < 0.10 in a 500kb window, collapsed to the most significant variant. The major histocompatibility complex (MHC) gene region was represented by one variant in the single most-significant LD block. Our first standardized analyses involved testing and evaluating within each dataset the associations between the schizophrenia polygenic risk score and each of the five substance use disorder diagnostic phenotypes: any substance use disorder, tobacco use disorder, alcohol use disorder, cannabis use disorder, and stimulant use disorder.

In each dataset and for each p-value threshold used for the PGC-SCZ2 results, the schizophrenia polygenic risk score was regressed against the five substance use disorder phenotypes using logistic regression in R (44). Age, sex, and the first ten population-stratification principal components were included as covariates.

The proportion of variance explained (R^2^) by the schizophrenia polygenic risk score was computed by comparing the regression model with the age, sex, and ten principal components to the regression model that includes the schizophrenia polygenic risk score variable in addition to age, sex and principal components. Because the analyses used logistic regression, the reported R^2^ is the difference between the Nagelkerke’s pseudo-R^2^ from the two models.

We then performed a mega-analysis where the individual level data were combined into one dataset and analyzed. We examined the association between the schizophrenia polygenic risk score and the five substance use disorder phenotypes (any substance use disorder diagnosis, tobacco use disorder, alcohol use disorder, cannabis use disorder, and stimulant use disorder), using a similar approach as described above, which now included an adjustment for study, as well as age, sex, and ten principal components.

## Results

We examined the association between polygenic risk score for schizophrenia, as defined using varying p-value cutoffs (p*T*) for any substance use disorder (i.e., having tobacco, alcohol, cannabis or stimulant use disorder, as defined above), and then for each specific substance: tobacco use disorder, alcohol use disorder, cannabis use disorder, and stimulant use disorder. Figure 2 shows the association between the schizophrenia polygenic risk scores and these phenotypes in each individual dataset and then as a mega-analysis.

**Figure 2:**
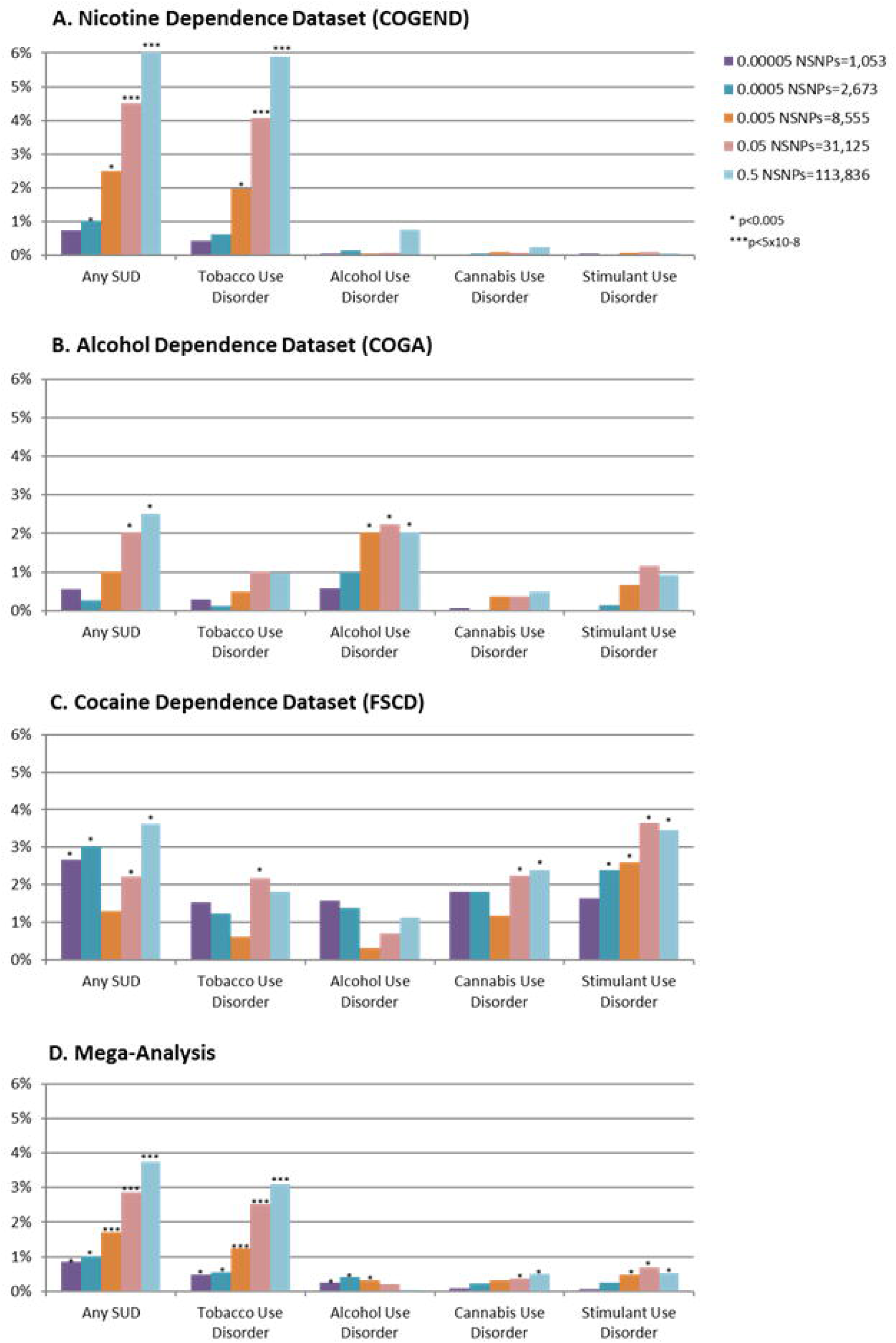
Polygenic risk scores for schizophrenia are associated with substance use disorder both in individual datasets (A-C) and in mega-analysis (D). *PT* is the p-value cutoff used to create the schizophrenia risk score, and the *R^2^* is from the regression between the substance-related phenotype and the schizophrenia risk score, adjusted for age, sex and principal components. In the mega-analyses, an additional adjustment for study ascertainment is included.

The phenotype most strongly and consistently associated with schizophrenia polygenic risk scores was the “any substance use disorder” phenotype. In general, the associations became stronger overall with less restrictive p-value cutoffs, which is consistent with previous studies (22, 23). In the mega-analysis, the association between any substance use disorder and schizophrenia polygenic risk score peaked at p*_T_*<0.5; pseudo R^2^ range 0.8% to 3.7%, minimum p = 4×10^-23^.

In each individual dataset, a statistically significant association was seen between polygenic risk score for schizophrenia and the substance of ascertainment. Specifically, tobacco use disorder was most strongly associated with schizophrenia polygenic risk score in COGEND, the dataset ascertained to study nicotine dependence (Figure 2A, pseudo R^2^ range 0.42%-5.8%, minimum p<9.5×10^-20^), alcohol use disorder was most strongly associated with schizophrenia polygenic risk score in COGA, the dataset ascertained to study alcohol dependence (Figure 2B, pseudo R^2^ range 0.57%-2.1%, minimum p<7×10^-6^), and stimulant use disorder was most strongly associated with schizophrenia polygenic risk score in FSCD, the dataset ascertained to study cocaine dependence (Figure 2C, pseudo R^2^ range 1.7-3.5%, minimum p<9×10^-5^). Cannabis use disorder was associated with polygenic risk score only in FSCD, the dataset ascertained to study cocaine dependence. These results led to statistically significant associations between schizophrenia polygenic risk score and tobacco, alcohol, cannabis, and stimulant use disorder in the mega-analysis (Figure 2D, tobacco use disorder pseudo R^2^ range 0.5%-3.1%, minimum p=2×10^-18^; alcohol use disorder pseudo R^2^ range 0.04%-0.4%, minimum p=4×10^-4^; cannabis use disorder pseudo R^2^ range 0.09%-0.5%, minimum p=0.001; stimulant use disorder pseudo R^2^ range 0.06%-0.7%, minimum p=5×10^-5^).

The most consistent association across all three datasets was between any substance use disorder and schizophrenia polygenic risk score, though a different primary substance use disorder contributed to this association depending on the ascertainment criteria for that dataset. In the mega-analysis, there was also a very strong association between tobacco use disorder and schizophrenia polygenic risk score. We suspect that this is in part because of the high level of comorbidity between tobacco use disorder and alcohol, cannabis, and stimulant use disorder.

The comorbidity among the substance use disorders is complex. First, cases from one study are much more likely than controls to have another comorbid substance dependence. For example, when nicotine use disorder is analyzed in the alcohol dataset, 48% of the non-nicotine dependent individuals have alcohol dependence and are included in the control group, and therefore the observed association is tempered. In order to evaluate the observed association between the schizophrenia polygenetic risk score and any substance use disorder diagnosis, we performed a secondary analysis in which we extracted individuals without tobacco use disorder from the combined dataset. This decreased the sample size in the mega-analysis from 3,488 to 1,657 participants. We repeated the testing of the association between polygenic risk score for schizophrenia (pT<0.5) and any substance use disorder, adjusted for study, age, sex, and principal components. Although the association remained statistically significant (p=0.0015), the adjusted odds ratio for the standardized score dropped from 1.55 (95% CI 1.42-1.69) to 1.25 (95% CI 1.08-1.44) and the proportion of variance explained dropped from 3.7% to 0.7%. Therefore, we cannot rule out the possibility that the observed association between any substance use disorder and schizophrenia polygenic risk score is primarily driven by tobacco use disorder.

## Discussion

These results provide strong systematic evidence of shared polygenic risk between schizophrenia and substance use disorder. Each independent substance use disorder dataset showed a strong shared genetic architecture with a genetic liability to schizophrenia, and the signal was greatly strengthened when all substance use disorders were considered together. However, because of comorbidity among the substance use disorder diagnoses, we cannot statistically test whether the observed association is driven by tobacco use disorder or a general substance use disorder liability. It is well known that substance use disorders are often comorbid with one another, and family and twin studies have demonstrated that the underlying genetic liability to substance use disorder has both common and specific genetic risk factors (46-49).

The importance of ascertainment is highlighted by these analyses. When we analyzed the association between the schizophrenia polygenic risk score and a specific substance use disorder diagnosis that was aligned with ascertainment, the associations were robust. Similarly, when the data were mega-analyzed comparing any substance use disorder versus no substance use disorder, a strong association with the schizophrenia polygenic risk scores was seen. In contrast, analyses in the individual datasets showed minimal or no statistically significant associations between specific substance use disorder diagnoses for non-ascertained substances and the polygenic risk score for schizophrenia.

These findings are consistent with a common underlying shared genetic liability to schizophrenia or substance use disorder. Other possible models for the observed association include mediation by schizophrenia and mediation by substance use disorder (Figure 1). The mediation by schizophrenia model is commonly referred to as a “self-medication” model of comorbidity, where the development of schizophrenia subsequently leads to the onset of substance use disorder under the hypothesis that individuals use and misuse a substance to reduce symptomatology. In this study, because the self-reported prevalence of psychotic symptoms in these data is less than 5%, the observed association between the polygenic risk score for schizophrenia and the substance use disorder phenotypes cannot be explained through mediation by schizophrenia. However, biological risk of schizophrenia may be expressed in ways other than psychotic symptoms, and mediation through subthreshold symptoms of schizophrenia could drive the association seen with substance use disorder. The next mechanism, the development of schizophrenia mediated by substance use disorder, cannot be directly tested in these datasets. In addition, because the majority of the schizophrenia datasets do not have substance use behaviors measured, a more comprehensive analysis of the contribution of polygenic variation related to schizophrenia versus substance use disorder cannot be undertaken at this time. However, as the better understanding of this shared liability is studied in the future, we may find that all three potential pathways play a role in the shared genetic liability between schizophrenia and substance use disorder.

Interestingly, in these data, the magnitudes of the phenotypic variance explained by schizophrenia polygenic risk score for substance use disorder are larger than other estimates of pseudo R^2^ for the association of polygenic risk score for schizophrenia and other phenotypes (12, 17, 50-52). However, it is important to note that R^2^ is an estimate specific to the individual datasets, and is difficult to extrapolate across studies. Nonetheless, we attribute the strong findings seen in these data to the sampling of phenotypic extremes of substance use disorder, where stringent ascertainment leads to a stronger model fit than previously reported (53).

Although the pseudo R^2^ estimates are unusually large, it highlights the statistical power in this sample for traits related to substance use disorder. It is common in genetic studies to meta-(and mega)-analyze the largest possible sample, which may combine studies with many different ascertainment schemes. Our results show that this approach of combining many different datasets with varying ascertainment schema may temper associations. For example, Chen et al. (22) found a much weaker association of nicotine dependence and cigarettes per day with a similarly generated polygenic risk score for schizophrenia in a large meta-analysis of samples ascertained as population and disease-based cohorts (R^2^ ≤ 0.1%, N=13,326). Our data suggest that when testing the relationship between a substance-related phenotype and the polygenic risk score for schizophrenia, the diagnosis of any substance use disorder may be the most informative substance-related phenotype to use, especially with a heterogeneously ascertained series of samples. Despite this, we would expect that analyses with large enough sample sizes, as in the study reported by Chen et al. (22), will detect attenuated associations between the schizophrenia polygenic risk score and individual substance use disorder. These findings highlight the power of carefully ascertained smaller samples where precise phenotyping can provide useful insights (and large effect sizes) that may not otherwise be seen.

Finally, the finding of shared genetic factors between substance use disorder and schizophrenia further challenges the diagnostic boundaries that typically separate substance use disorder from both psychotic and mood disorders. Since the birth of DSM in 1952 (54), there has been a sharp distinction between substance use disorder and psychotic disorders. Increasing, scientific evidence supports a blurred biological boundary and will hopefully lead to improved understanding of the neurophysiology underlying both disorders.

## Acknowledgements

Funding for this study was provided by the National Institutes of Health (K08DA032680, R21AA024888, R01DA035825, R01DA036583, K02DA32573, R01DA023668, R01AG045231, U01AG052564, and R01HD083614), and the National Science Foundation (DGE-1143954).

Funding support for the CIDR-COGA Study was provided through the Center for Inherited Disease Research (CIDR) and the Collaborative Study on the Genetics of Alcoholism (COGA). Support for collection of datasets and samples and assistance with phenotype harmonization and genotype cleaning, as well as with general study coordination, was provided by the Collaborative Study on the Genetics of Alcoholism (COGA; U10AA008401). Assistance with data cleaning was provided by the National Center for Biotechnology Information. Funding support for genotyping, which was performed at the Johns Hopkins University Center for Inherited Disease Research, was provided by the NIH GEI (U01HG004438), the National Institute on Alcohol Abuse and Alcoholism, and the NIH contract “High throughput genotyping for studying the genetic contributions to human disease” (HHSN268200782096C). The datasets used for the analyses described in this manuscript were obtained from dbGaP at http://www.ncbi.nlm.nih.gov/gap through dbGaP accession number phs000125.v1.p1.

The Family Study on Cocaine Dependence was genotyped as part of the Study of Addiction: Genetics and Environment (SAGE). Funding support for SAGE was provided through the NIH Genes, Environment and Health Initiative [GEI] (U01 HG004422). SAGE is one of the genome-wide association studies funded as part of the Gene Environment Association Studies (GENEVA) under GEI. Assistance with phenotype harmonization and genotype cleaning, as well as with general study coordination, was provided by the GENEVA Coordinating Center (U01 HG004446). Assistance with data cleaning was provided by the National Center for Biotechnology Information. Support for collection of datasets and samples was provided by the Collaborative Study on the Genetics of Alcoholism (COGA; U10 AA008401), the Collaborative Genetic Study of Nicotine Dependence (COGEND; P01 CA089392), and the Family Study of Cocaine Dependence (FSCD; R01 DA013423, R01 DA019963). Funding support for genotyping, which was performed at the Johns Hopkins University Center for Inherited Disease Research, was provided by the NIH GEI (U01HG004438), the National Institute on Alcohol Abuse and Alcoholism, the National Institute on Drug Abuse, and the NIH contract “High throughput genotyping for studying the genetic contributions to human disease” (HHSN268200782096C). The datasets used for the analyses described in this manuscript were obtained from dbGaP at http://www.ncbi.nlm.nih.gov/gap through dbGaP accession number phs000092.v1.p1.

### Conflict of interest

The authors have no competing financial interest in relation to the work described.

